# FUME-TCRseq: Sensitive and accurate sequencing of the T-cell receptor from limited input of degraded RNA

**DOI:** 10.1101/2023.04.24.538037

**Authors:** Ann-Marie Baker, Gayathri Nageswaran, Pablo Nenclares, Tahel Ronel, Kane Smith, Christopher Kimberley, Miangela M Lacle, Shree Bhide, Kevin J Harrington, Alan Melcher, Manuel Rodriguez-Justo, Benny Chain, Trevor A Graham

**Affiliations:** Centre for Evolution and Cancer, Institute of Cancer Research, London, United Kingdom; Centre for Genomics and Computational Biology, Barts Cancer Institute, Queen Mary University of London, London, United Kingdom; Division of Infection and Immunity, University College London, London, United Kingdom; Division of Radiotherapy and Imaging, The Institute of Cancer Research, London, United Kingdom; Department of Pathology, University Medical Center Utrecht, Utrecht, Netherlands; Department of Histopathology, UCL Hospitals NHS Trust, London, UK

**Author notes:** denotes co-senior authors (Benny Chain, Trevor A Graham). **Author contributions** A-MB and PN performed wet-lab experiments and data analysis, with computational support from TR and experimental support from GN, KS and CK. MML, MR-J, SB and KH identified patient samples, with MML and MR-J providing expert histological guidance. Study conception and design was by A-MB, BC and TAG, and study supervision was by AM, BC and TAG. A-MB, BC and TAG wrote the first draft of the manuscript, with all authors editing and approving the final version.

## Abstract

Genomic analysis of the T-cell receptor (TCR) reveals the strength, breadth and clonal dynamics of the adaptive immune response to pathogens or cancer. The diversity of the TCR repertoire, however, means that sequencing is technically challenging, particularly for samples with low quality, degraded nucleic acids. Here, we have developed and validated FUME-TCRseq, a robust and sensitive RNA-based TCR sequencing methodology that is suitable for formalin-fixed paraffin-embedded samples and low amounts of input material. FUME-TCRseq incorporates unique molecular identifiers into each molecule of cDNA, allowing correction for sequencing errors and PCR bias. We used RNA extracted from colorectal and head and neck cancers to benchmark the accuracy and sensitivity of FUME-TCRseq against existing methods, and found excellent concordance between the datasets. Furthermore, FUME-TCRseq detected more clonotypes than a commercial RNA-based alternative, with shorter library preparation time and significantly lower cost. The high sensitivity and the ability to sequence RNA of poor quality and limited amount enables quantitative analysis of small numbers of cells from archival tissue sections, which is not possible with other methods. To demonstrate this we performed spatially-resolved FUME-TCRseq of colorectal cancers using macrodissected archival samples, revealing the shifting T-cell landscapes at the transition to an invasive phenotype, and between tumour subclones containing distinct driver alterations. In summary, FUME-TCRseq represents an accurate, sensitive and low-cost tool for the characterisation of T-cell repertoires, particularly in samples with low quality RNA that have not been accessible using existing methodology.

## Introduction

T cells are critical drivers of the adaptive immune response, whose antigen specificity is determined by the highly diverse T-cell receptor (TCR) sequence. High throughput sequencing and analysis of the TCR repertoire have emerged as powerful tools for profiling T-cell responses to pathogens or cancer, or indeed to host tissues in autoimmune disease, however sequencing the TCR locus is particularly challenging. In this manuscript we describe the development, validation, and application of a new TCR sequencing (TCRseq) methodology.

The TCR is a highly diverse heterodimer that is expressed on the surface of T cells. The majority of T cells express an αβ TCR, whereas γδ T cells generally represent a smaller proportion of the T-cell population. The loci of TCR chains are arranged in segments, with a variable (V) region, a diversity (D) region (β and δ only) and a joining (J) region, followed by a constant (C) region. The TCR is produced by a series of stochastic DNA recombination processes which occur in the early stages of T cell differentiation in the thymus. During T-cell maturation one allele of each segment will randomly recombine with the others to form a functional TCR. The β and δ chains undergo VDJ recombination, whereas the α and γ chains undergo VJ recombination. In addition there is random nucleotide insertion and deletion at the junctions between recombined segments, generating further diversity. The most variable region of the TCR is the complementarity determining region 3 (CDR3), which is a key component of antigen specificity. The chance of independent generation of the same TCR sequence in a single individual is incredibly low, therefore sequencing the CDR3 region of the TCRβ gene can be used as a unique identifier of a T-cell clone.

The processes that contribute to the creation of a highly diverse TCR repertoire pose formidable challenges to its genetic analysis. Firstly, since DNA recombination is somatic, the TCRβ locus cannot be predicted from the germline sequence. Secondly, the stochastic nature of DNA recombination which results in an enormous amount of genomic diversity means that sequencing across the TCRβ region is particularly challenging. Finally, since clonal amplification of T cells is a fundamental aspect of an immune response, it becomes crucial not only to sequence the TCRs produced in all T cells but to do so quantitatively.

Library preparation for TCR sequencing can use either genomic DNA (gDNA) or total RNA as input material, and these approaches have been benchmarked^1,2^ and reviewed^3^ elsewhere. Protocols that use gDNA have the advantage of a 1:1 relationship between number of cells and sequence abundance, however non-expressed, potentially irrelevant sequences will be detected^3^. In contrast, protocols that use RNA have more functional relevance as only expressed transcripts are sequenced, and furthermore they require less input and are more sensitive to rare clonotypes. However, RNA is less stable than DNA, and therefore many commercially available protocols require high-quality RNA as input. Indeed, the majority of existing TCR sequencing methods are not effective for poor-quality input material, such as highly degraded RNA extracted from formalin-fixed paraffin-embedded (FFPE) samples. The few commercial kits that claim to be FFPE-compatible have displayed a poor success rate in our hands and are prohibitively expensive for many researchers. Therefore, we recognised an urgent need for a robust, low-cost and FFPE-compatible RNA-based TCR sequencing methodology.

Here we describe our novel protocol, FUME-TCRseq (FFPE-suitable Unique Molecular idEntifier-based TCRseq). The method is multiplex PCR-based, and uses 38 primers against the TCRB V genes. Crucially it incorporates unique molecular identifiers (UMIs) for the correction of amplification bias and sequencing errors. UMIs are essential for quantitative and robust analysis of TCR repertoire data and are not included in many of the current TCRseq methods. Compared to TCRseq methods that do include UMIs, the novelty of FUME-TCRseq is that the UMI is on the reverse transcription primer; the method does not involve template switching or second strand synthesis. We show that omitting these inefficient steps improves the sensitivity of TCR clonotype detection. Furthermore, FUME-TCRseq targets an amplicon of approximately 170 base pairs, which is shorter than other multiplex methods and enables analysis of highly degraded RNA. FUME-TCRseq requires no specialist equipment and costs less than £30 per sample (approximately 10-fold cheaper than commercial equivalents), meaning that it will be accessible to most genomic and molecular biology laboratories.

## Results

### FUME-TCRseq data is consistent with 5’-RACE TCRseq

FUME-TCRseq is a novel protocol that combines UMI incorporation and enrichment of the TCRβ CDR3 region using multiplex PCR (Figure 1). We limit our analysis to the hypervariable CDR3 region of TCRβ as it is responsible for the majority of antigen specificity. Our UMIs are 12 random base pairs (two sets of 6 base pairs, separated by an 8 base pair spacer). The addition of UMIs at reverse transcription means that sequencing reads with the same UMI can be traced back to the same initial strand of cDNA. The first advantage of using UMIs is it enables the correction of PCR and sequencing errors. This is particularly important for TCR sequencing, as TCR repertoires contain non-template sequences which can differ by only one or a few nucleotides from germline sequences and from each other, so it is crucial to determine which sequences represent true T-cell rearrangements and which are artificially generated by errors in PCR or sequencing. Second, the use of UMIs can correct for inherent bias in PCR amplification, allowing more accurate quantification of the relative abundance of each T cell clone. After reverse transcription we amplify TCRβ CDR3 using a pool of 38 V gene primers (Supplementary Table 1) that were developed and optimised by the EuroClonality-NGS consortium for the monitoring of lymphoid malignancies^4^.

**Figure 1.**
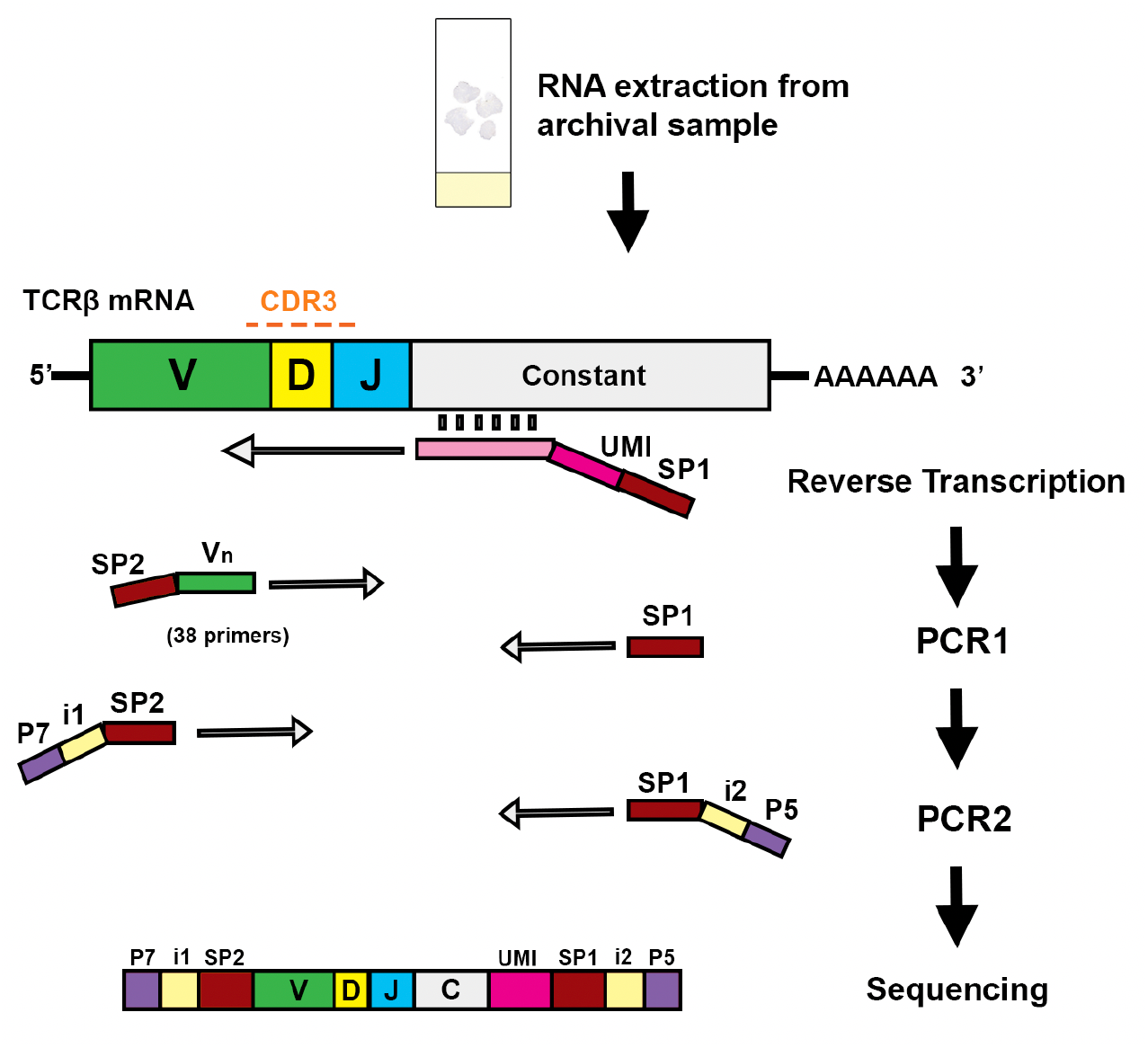
Schematic representation of the FUME-TCRseq methodology, whereby RNA is reverse transcribed using a UMI-containing primer, followed by multiplex PCR amplification of the CDR3 region using a panel of 38 primers, and a second PCR to add sample-specific indexes.

As a first validation of FUME-TCRseq, we sequenced high-quality RNA extracted from a sample of whole blood. In parallel, we used a well-established 5’-RACE ligation-based method^5^ using the same pool of RNA as input. Notably, FUME-TCRseq produced a higher proportion of reads mapping to the TCR than the 5’RACE method (98.0% vs 61.4%, Supplementary Table 2). FUME-TCRseq data was randomly downsampled to the same number of decombined (“mapped”) reads as the 5’-RACE data (538942 reads). FUME-TCRseq called many more clonotypes than the ligation method (113981 vs 15512, Supplementary Table 2), and interestingly had a lower proportion of non-productive sequences (4.1% vs 15.5%, Supplementary Table 2). We found that the data on V gene usage were highly consistent between the two methods (Figure 2, A and B), indicating that the multiplex PCR approach does not skew the data in favour of particular V genes. We next examined the data for overlap between the specific clonotypes detected. The overlap between the methods was 2803 clonotypes (18.1% of the ligation method repertoire, 2.5% of the FUME-TCRseq repertoire). However when we considered only the most expanded clonotypes (those represented by more than 5 UMIs in the ligation method data), this increased to 74/78 (94.9%). These results indicates a high level of sampling effect in the data. Conversely, the most common clonotypes detected by FUME-TCRseq are also detected by the ligation method, and at similar relative frequency (Figure 2C).

**Figure 2.**
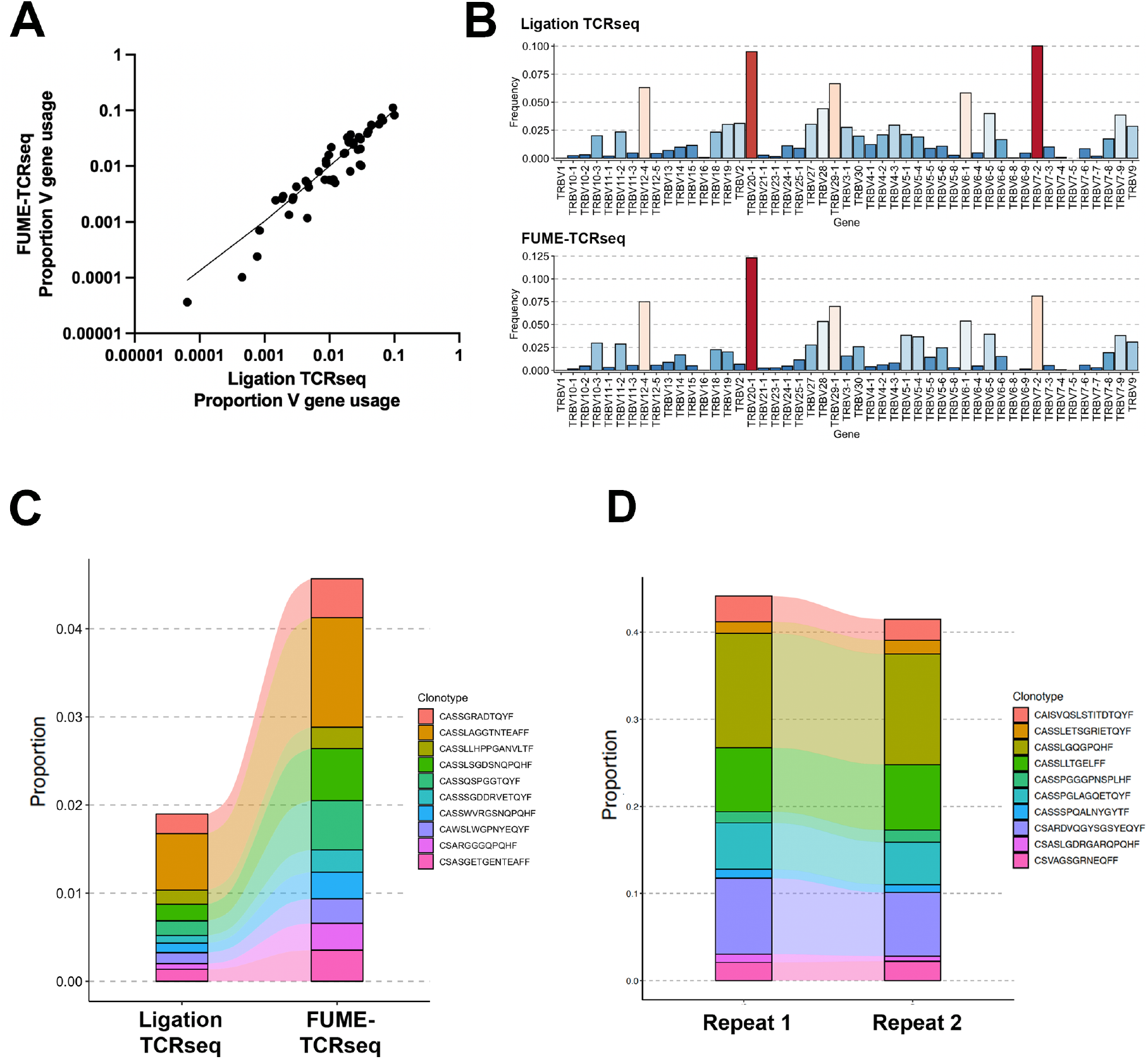
**A**. Scatter plot showing V gene usage data from FUME-TCRseq and 5’-RACE ligation methodologies, performed on RNA extracted from a representative whole blood sample. **B**. Bar plot showing the proportion of each V gene detected by FUME-TCRseq and 5’-RACE ligation. Red bars are highly frequent V genes, and blue are less frequent. **C**. Plot showing the 10 most common clonotypes identified by FUME-TCRseq, and the frequencies at which they were detected using 5’-RACE ligation methodology. **D**. Plot showing the 10 most common clonotypes detected in technical repeats of FUME-TCRseq performed on RNA extracted from an inflammatory polyp.

We next extracted high quality RNA from a fresh-frozen biopsy of a polyp from a patient with inflammatory bowel disease, anticipating a diverse immune repertoire likely with expansions of specific T-cell clonotypes. We tested the reproducibility of FUME-TCRseq by performing duplicate library preparations using the same pool of input RNA. A similar number of unique productive TCR sequences was detected in the replicates (1,787 vs 2,100). Overall, we detected 34.7% (620/1787) of the clonotypes on repeat, but 98.6% (71/72) of the most abundant clones (represented by more than 20 UMIs). Importantly, we detected the most expanded clonotypes in both repeats at very similar proportions (Figure 2D), confirming good reproducibility of FUME-TCRseq.

### FUME-TCRseq identifies novel T cell clonotypes in tumour subclones

In our recent multi-omic analysis of primary colorectal cancers we identified two tumours with subclonal mutations in driver genes^6^. We applied the BaseScope assay (point mutation-specific RNA *in situ* hybridisation^7^) to an FFPE tissue section to precisely identify cells expressing mutant transcripts (Figure 3A). Using this annotation as a guide we macrodissected wild-type (WT) and mutant regions of similar size from serial, unstained FFPE tissue sections of these tumours. We then extracted RNA from these regions and performed FUME-TCRseq. We note that these RNA samples were highly degraded, with RNA integrity number (RIN) of 1.0 (C537) and 2.3 (C539).

**Figure 3.**
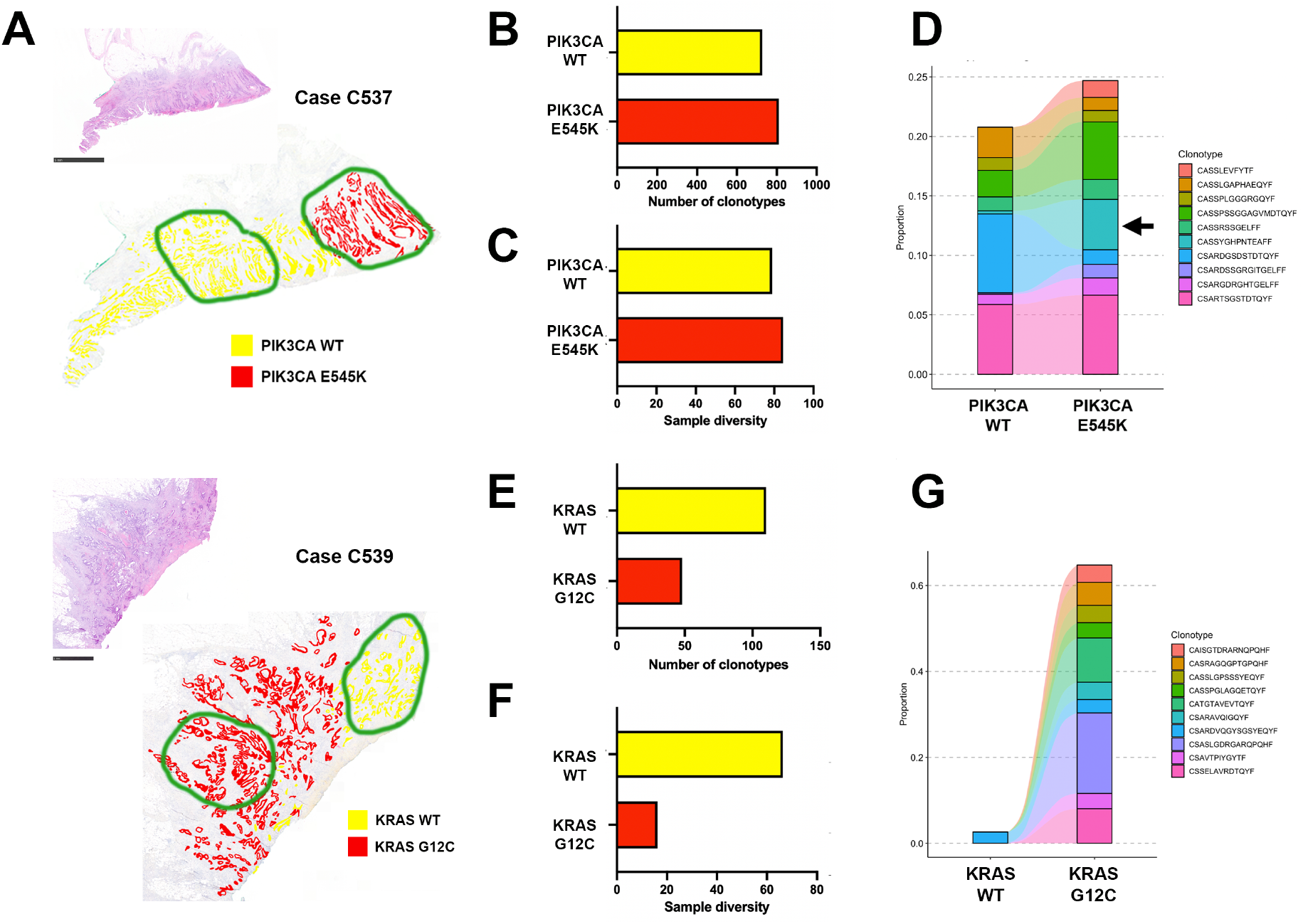
**A**. H&E images and annotated BaseScope staining for point mutations in PIK3CA (Case C537, upper panel) or KRAS (Case 539, lower panel). Scale bars represent 5mm (upper panel) and 1mm (lower panel). Regions circled in green were macrodissected for FUME-TCRseq. **B**. Bar chart showing the numbers of unique TCR clonotypes detected in the PIK3CA wild-type (WT) and mutant (E545K) subclones of Case C537. **C**. Bar chart showing the diversity of the TCR repertoire in PIK3CA wild-type and mutant subclones of Case C537. **D**. Plot showing the frequencies of the 10 most common clonotypes detected in the PIK3CA E545K mutant subclone, and the frequencies at which they appear in the wild-type subclone. Indicated with an arrow is the TCR sequence CASSYGHPNTEAFF. **E**. Bar chart showing the numbers of unique TCR clonotypes detected in the KRAS wild-type (WT) and mutant (G12C) subclones of Case C539. **F**. Bar chart showing the diversity of the TCR repertoire in KRAS wild-type and mutant subclones of Case C539. **G**. Plot showing the frequencies of the 10 most common clonotypes detected in the KRAS G12C mutant subclone, and the frequencies at which they appear in the wild-type subclone.

In case C537 we found that the PIK3CA wild-type and E545K-mutant subclones had a similar number of unique TCR clonotypes (727 vs 810, Figure 3B, Supplementary Table 2) and similar diversity by the Inverse Simpson index (78.9 vs 84.5, Figure 3C). We then compared the ten most common TCR clonotypes in the WT and mutant regions and found that although there were no expanded clonotypes unique to the WT region, there were three clonotypes expanded only in the mutant subclone. Of particular interest is the CDR3 sequence “CASSYGHPNTEAFF” which represented 3.5% of all TCR reads in the PIK3CA E545K-mutant subclone, and only 0.21% in the PIK3CA WT subclone (Figure 3D).

Furthermore, this CDR3 amino acid sequence was detected six times in the mutant repertoire, with different nucleotide sequences, indicating T-cell convergence towards this clonotype. The spatial distribution of the TCR sequence is suggestive of specificity to a newly generated subclone-specific neoantigen, although we note that this neoantigen is not necessarily the PIK3CA E545K mutation itself.

The subclonal TCR repertoires of case C539 were strikingly different. The KRAS G12C mutant subclone had considerably fewer unique productive TCR clonotypes than the WT subclone (48 vs 110, Figure 3E, Supplementary Table 2) and lower diversity by the Inverse Simpson index (16.2 vs 66.3, Figure 3F). We noted that the WT subclone had 1.7x more sequencing reads than the mutant, therefore we randomly downsampled the WT mapped reads to match that of the mutant and repeated our analysis (Supplementary Table 2). This did not drastically change the number of unique clonotypes (all data = 110, downsampled data = 108) or the diversity (all data = 66.3, downsampled data = 68.8). When we examined the ten most expanded clonotypes in the KRAS G12C mutant subclone, only one was present in the KRAS WT subclone (Figure 3G). These results indicate a lesser T-cell response to the KRAS G12C mutant subclone.

### FUME-TCRseq reveals an altered TCR repertoire at the transition to invasion

In a recent study we used mathematical modelling combined with sequencing and image-based analysis of the microenvironment to infer the presence of an immune bottleneck at the transition from a benign colorectal adenoma to invasive carcinoma^8^. Here we sought to characterise the T-cell repertoires at the invasive transition by performing TCRseq on macrodissected matched adenoma and carcinoma regions (of similar size) from a single FFPE tissue section (Figure 4A). RNA extracted from these regions had a RIN of 2.3.

**Figure 4.**
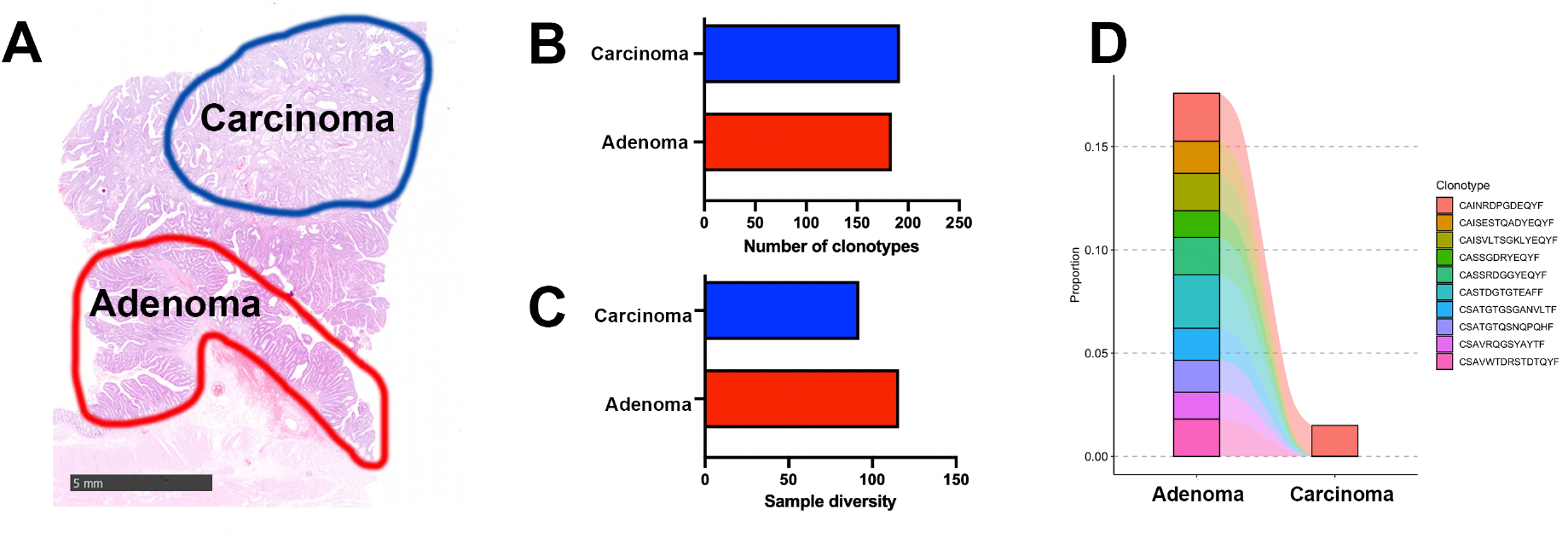
**A**. H&E showing the regions of FFPE adenoma and carcinoma that were macrodissected for RNA extraction and FUME-TCRseq. Scale bar represents 5mm. **B**. Bar chart showing the numbers of unique TCR clonotypes detected in carcinoma and adenoma regions. **C**. Bar chart showing the diversity of the TCR repertoire in carcinoma and adenoma regions. **D**. Plot showing the frequencies of the 10 most common clonotypes detected in the adenoma region, and the frequencies at which they appear in the carcinoma.

We found that the number of unique productive TCR clonotypes detected in adenoma and carcinoma regions was broadly similar (184 vs 192, Figure 4B, Supplementary Table 2); however the carcinoma TCR repertoire was less diverse by the Inverse Simpson index (92.2 vs 116.0, Figure 4C). This indicates a broader immune response in the adenoma, with the TCR repertoire of the carcinoma being more focused. The regions share only one clonotype in their ten most expanded clonotypes (Figure 4D), suggesting a very different T-cell response to cells of the adenoma and carcinoma.

### Comparison of FUME-TCRseq data with industry standards

We compared the performance of FUME-TCRseq against two commercially available methods. First, we chose the Immunoverse assay (ArcherDx), as it uses RNA as input material and is a multiplex PCR-based protocol that incorporates UMIs. To our knowledge it is the most similar protocol to FUME-TCRseq that is currently commercially available, therefore we considered it to be the most appropriate comparator.

We used the WT and mutant samples from C537 (Figure 3A-D) to compare the methods, running both protocols in parallel with the same amount of input RNA from the same RNA pool. We looked at V gene usage and found it to be broadly consistent between the methods (Figure 5A), with TRBV20-1 the most frequently used V gene in all cases, representing between 12.7-19.7% of all clonotypes. The proportions were lower in the Immunoverse samples, likely because of the large number of “ambiguous” V gene calls.

**Figure 5.**
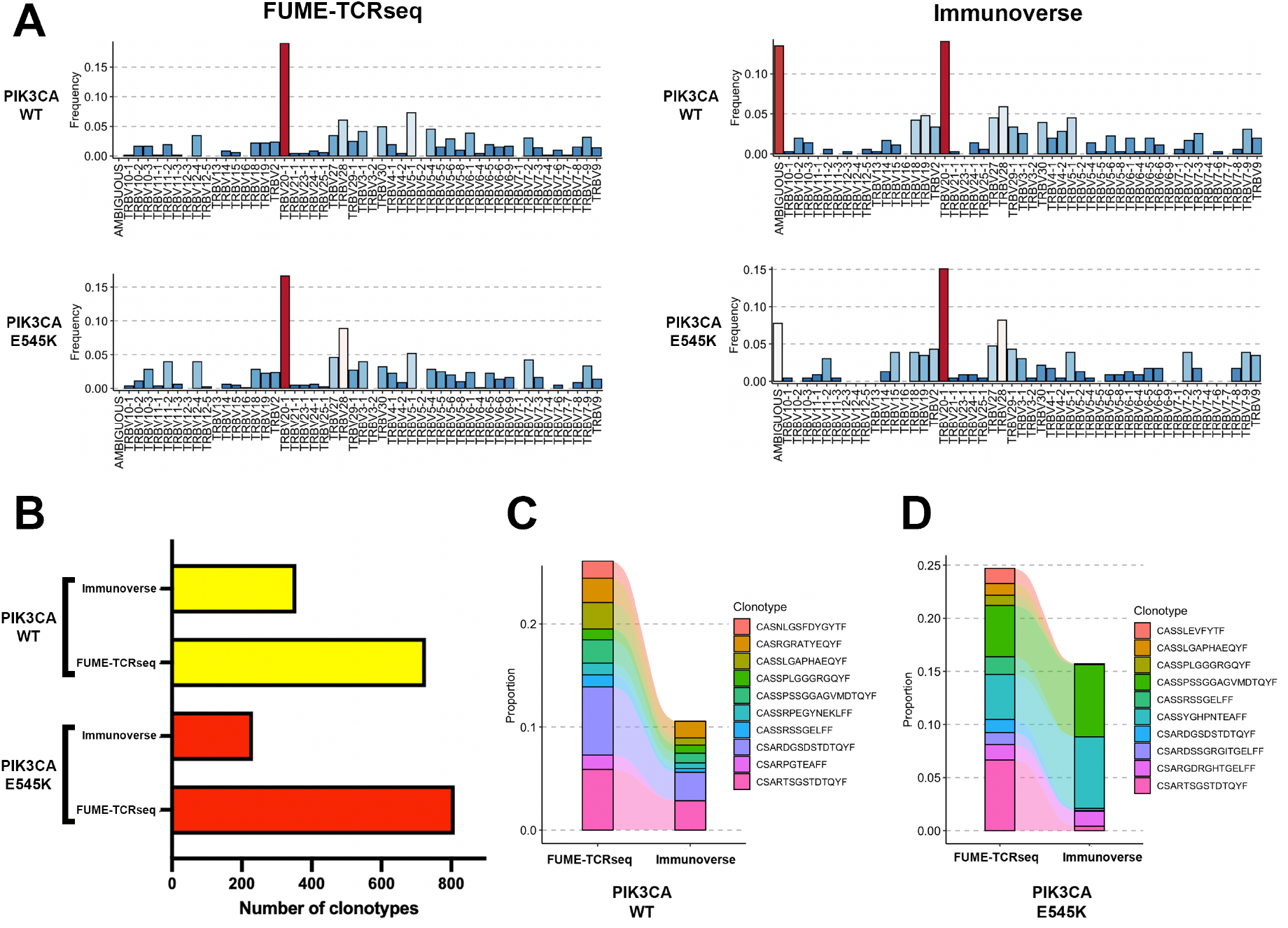
**A**. Bar plots showing the proportion of each V gene detected by FUME-TCRseq (left column) and Immunoverse (right column) in two representative FFPE colorectal cancer samples (“PIK3CA WT”-top row, “PIK3CA E545K” – bottom row). **B**. Bar chart showing the numbers of unique TCR clonotypes detected in the two samples with FUME-TCRseq and Immunoverse methodologies. **C**. Plot showing the frequencies of the 10 most common clonotypes detected in the wild-type sample by FUME-TCRseq, and the frequencies at which they appear in the Immunoverse data. **D**. Plot showing the frequencies of the 10 most common clonotypes detected in E545K sample by FUME-TCRseq, and the frequencies at which they appear in the Immunoverse data.

In both WT and mutant samples, the Immunoverse assay detected fewer unique clonotypes than FUME-TCRseq (Figure 5B, 356 vs 727 for wild-type region, 232 vs 810 for mutant region, Supplementary Table 2). Although a similar number of sequencing reads per sample was generated for both methods, Immunoverse results had only around 10% on-target reads, whereas FUME-TCRseq had around 70%. This is likely to be a contributing factor to the lower number of clonotypes detected by Immunoverse related to FUME-TCRseq.

We next examined the concordance between the clonotypes detected in the methods. Of 727 clonotypes detected by FUME-TCRseq of sample C537 WT, 105 (14.4%) of these were detected by the Immunoverse assay. Considering only the ten most expanded clones, eight of these were detected by Immunoverse (Figure 5C). Of the 810 clonotypes detected in FUME-TCRseq of sample C537 mutant, there were 62 (7.7%) detected by Immunoverse. Of the ten most expanded clones detected by FUME-TCRseq, seven were detected by Immunoverse (Figure 5D). Interestingly, the previously highlighted clone of interest identified by comparing the WT to mutant regions (“CASSYGHPNTEAFF”) was detected by Immunoverse, and at a similarly high proportion of the total TCR repertoire (6.3% vs 3.5%).

We next compared FUME-TCRseq to the immunoSEQ assay (Adaptive Biotechnologies), which uses gDNA as input material. We extracted DNA and RNA from the same macrodissected tissue isolated from three FFPE head and neck tumours, and used the DNA for immunoSEQ and the RNA for FUME-TCRseq. There were notable differences in V gene usage between the methods (Figure 6A), which was expected as immunoSEQ detects sequences that are not transcribed and is therefore subject to background amplification of non-functional chains. In cases HN1 and HN3, FUME-TCRseq detected fewer clonotypes than immunoSEQ, and in case HN2 it detected slightly more (Figure 6B, Supplementary Table 2); however the Morisita overlap between paired immunoSEQ and FUME-TCRseq repertoires was high (Figure 6C). Of all the clonotypes detected by FUME-TCRseq, a mean of 27.2% were detected in the paired immunoSEQ sample. The majority of the most expanded clones detected by ImmunoSEQ were also detected by FUME-TCRseq (Figure 6D), although relative abundances often varied.

**Figure 6.**
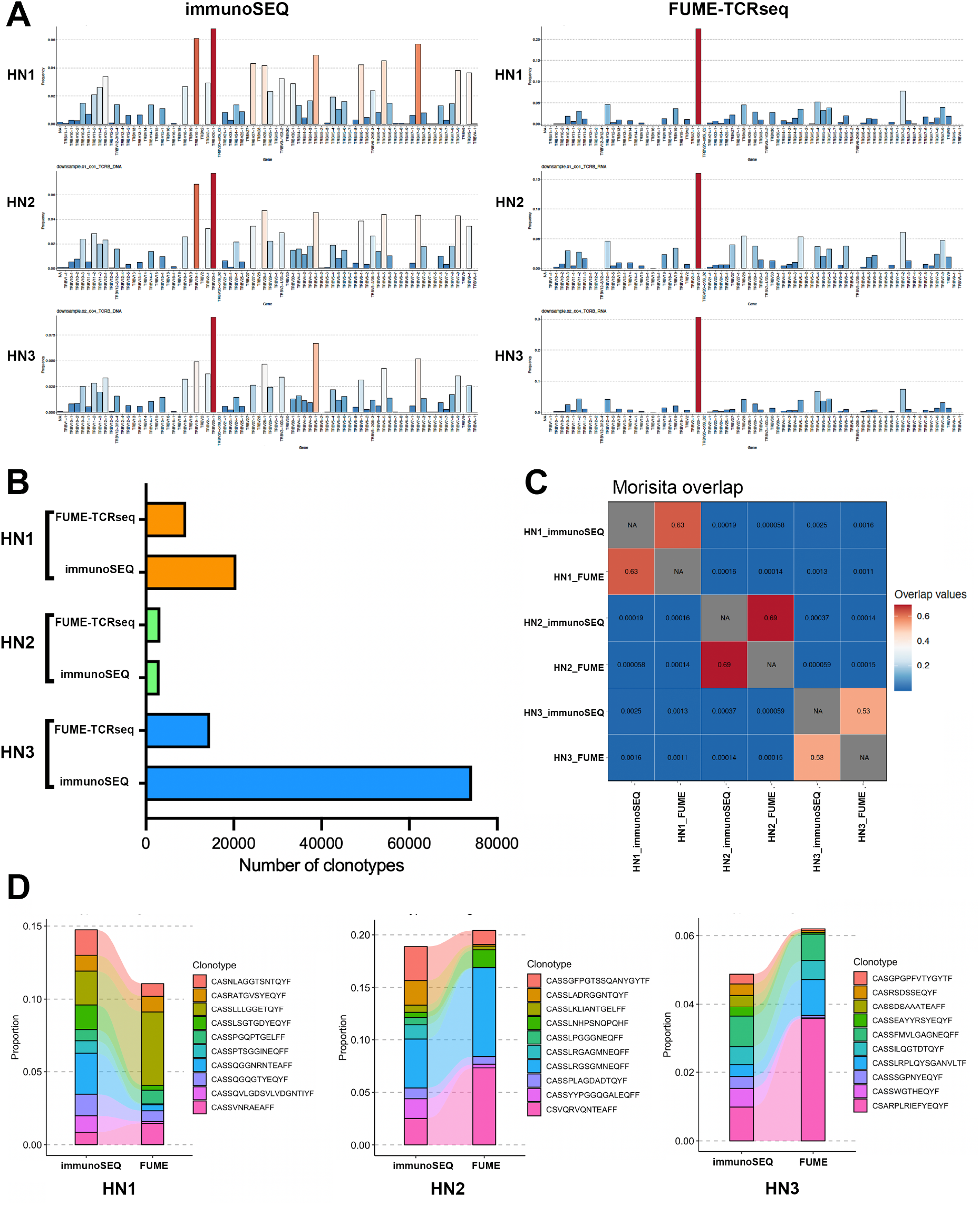
**A**. Bar plots showing the proportion of each V gene detected by FUME-TCRseq (right column) and immunoSEQ (left column) in three representative FFPE head and neck cancer samples (HN1, HN2, HN3). **B**. Bar chart showing the numbers of unique TCR clonotypes detected in the three samples with FUME-TCRseq and immunoSEQ methodologies. **C**. Heatmap showing the Morisita overlap between each sample. **D**. Plots showing the frequencies of the 10 most common clonotypes detected in each sample by immunoSEQ, and the frequencies at which they appear in the FUME-TCRseq data.

## Discussion

Profiling of the T-cell receptor repertoire has wide-ranging applications in health and disease; however the majority of existing methodologies require large amounts of high-quality input material. Therefore, to date, TCR repertoire analysis of archival FFPE tissue has not been feasible. This manuscript describes FUME-TCRseq, a novel assay for TCR repertoire profiling in highly degraded samples. It is the FFPE-compatibility and error correction via incorporation of UMIs which represent a significant methodological advance over other multiplex PCR-based approaches^9^.

We validated the protocol by comparing FUME-TCRseq to a well-established RACE-based TCRseq protocol^5^. We found no bias in the detection of V genes, and revealed a high rate of concordance in the detection of specific clonotypes, particularly when only expanded clones were considered. Furthermore, we benchmarked our protocol against a FFPE-compatible commercial kit (Immunoverse, ArcherDX) and found that our method detected a higher number of TCR clonotypes. Moreover, when comparing our protocol with a commercially available DNA-based TCRseq methodology (ImmunoSEQ) we found that our method detected a smaller number of unique TCRs, most likely due to the fact that DNA-based methodologies tend to overestimate T-cell divergence^1^. However, the pair-wise repertoire overlap by the Morisita index was high.

The sensitivity of our method is conducive to novel spatially-resolved TCRseq from macrodissected FFPE material, thus enabling multiregion profiling of the T-cell repertoire and correlation with histological features. We exemplified this by profiling distinct genetic and morphological subclones of primary colorectal cancers and revealing new insights into immune system co-evolution with tumour cells.

FUME-TCRseq is highly successful in FFPE archival samples, even those with RIN score less than 2. However, we note that FUME-TCRseq library preparation can fail for some samples, and this seems to be largely attributable to particularly low T-cell content. A further limitation of our method is that we only sequence the CDR3 region of the β chain, although this is often considered to be a good surrogate for T-cell clonal identity. Extension of the method to α chain is currently under exploration.

Although we developed FUME-TCRseq for the analysis of highly degraded samples, we have found the protocol to be very robust in the analysis of high-quality RNA samples as well. Therefore, it can be considered a universally applicable method, which is accessible to most researchers due to its low cost and ease of implementation.

In conclusion, FUME-TCRseq is a robust and sensitive novel method for T-cell repertoire profiling, with particular application to highly degraded samples that have hitherto been inaccessible. We anticipate that unlocking the analysis of archival FFPE tissue will facilitate longitudinal analysis of T-cell dynamics in clinical samples, and this has particular relevance in tracking immune responses through disease course and treatment. Broadly, this could reveal novel avenues for biomarker identification and drug development.

## Materials and Methods

### Sample collection

The sample of whole blood was collected and sequenced as part of a previously published study^10^. The fresh-frozen sample from a patient with inflammatory bowel disease was obtained from St Marks Hospital London, under Research Ethics Committee approval 18/LO/2051, with the patient giving informed consent. Formalin-fixed paraffin-embedded (FFPE) colorectal cancer samples were collected by the UCLH Cancer Biobank (Research Ethics Committee approval 15/YH/0311) and the University Medical Center Utrecht. The diagnostic FFPE head and neck cancer biopsies were obtained from The Royal Marsden Hospital, under the INSIGHT-2 study (CCR4934). Institutional board and ethics committee (ref. no. 19/LO/0638) approved the study.

### RNA extraction

RNA was extracted from the whole blood sample as previously described^10^. RNA from macrodissected histological tissue sections was extracted using the Roche HighPure FFPET RNA isolation kit (for FFPE samples) or the AllPrep DNA/RNA mini kit (for fresh frozen samples). RNA was quantified using the Qubit 3.0 fluorometer (Thermo Fisher), and RNA integrity number (RIN) was measured using the Agilent Tapestation 4200.

### RNA/DNA co-extraction from FFPE head and neck tumour samples

Five 10μm unstained slides and one hematoxylin and eosin-stained slides were obtained from representative FFPE tumour blocks. Experienced pathologists assessed tumour content and suitable areas of tumour were marked for macrodissection, if necessary. RNA and DNA were extracted using the AllPrep DNA/RNA FFPE kit (Qiagen). Nucleic acid yield and quality were assessed as described above.

### FUME-TCRseq library preparation and sequencing

A schematic of the FUME-TCRseq protocol is given in Figure 1, and we provide detailed descriptions of each step below.

#### Step 1. DNase treatment

After quantification, RNA is resuspended in RNase free water for a final volume of 8 μL. We recommend an input of 50ng for high quality RNA (RIN ≥ 7) or 100ng for samples with low integrity RNA (RIN < 7). Samples undergo DNase treatment by mixing 8 μL RNA, 1 μL RQ1 DNase (Promega), 1 μL RQ1 10× Buffer for 30 min at 37°C, then after addition of 1 μL RQ1 DNase stop buffer samples are incubated at 65°C for 10 minutes for inactivation.

#### Step 2. Reverse transcription

Firstly, 6.25 μL of RNase-free water, 1.5 dNTPs (10 mM each) and 0.75 μL “RT oligo” (10 μM, see Supplementary Table 1) are added to the 11 μL DNase treated RNA, and incubated at 65°C for 5 minutes, before immediately placing on ice for at least 1 minute. Next, 1.5 μL Superscript reverse transcriptase (Invitrogen ThermoFisher), 1.5 μL RNasin (Promega), 1.5 μL dithiothreitol (0.1 M), and 6 μL 5× Superscript IV buffer are added, and the samples are incubated at 55°C for 20 minutes, then 80°C for 10 minutes.

#### Step 3. Cleanup of cDNA

This purification and all subsequent purification steps are carried out using CleanNGS bead (GC Biotech) according to manufacturer’s instructions. The 30 μL reverse transcription product is mixed with 27 μL (0.9x) of CleanNGS beads. After incubation for 5 minutes at room temperature the beads are collected by placing on a magnetic stand for 2 minutes. The liquid above the beads in carefully aspirated and discarded and the beads are washed twice with 200 μL of freshly prepared 80% ethanol (EtOH), air-dried for 5 minutes and the DNA is eluted in 17.5 μL nuclease-free water. After placing on the magnetic stand for 2 minutes, 16.75 μL of cDNA is transferred to a new tube for PCR1.

#### Step 4. PCR1

A multiplex PCR using the oligo SP1 and a pool of 38 primers designed against the V regions of TCRb (see Supplementary Table 1) is performed. The V primer pool has been previously described^4^. 16.75 μL of cDNA is mixed with dNTPs (0.5 μL, 10 mM stock), SP1 primers (1.25 μL, 10 μM stock), V beta primer pool (1.25 μL), Phusion High Fidelity proofreading DNA polymerase (0.25 μL Thermofisher), and Phusion HF buffer (5 μL, 5 × stock). Initial denaturation was at 98°C for 3min, followed by PCR cycles (98°C 15sec, 54°C 30 sec, 72°C 30sec), and final elongation at 72°C for 5mins. For high quality RNA input (from PBMCs or fresh frozen tissue) 10 cycles of PCR was used, and for low quality (FFPE samples) 13 cycles was used.

#### Step 5. Cleanup of PCR1

A second purification step using CleanNGS beads at 0.75x (18.75 μL) is performed as previously described in Step 3. The purified product is eluted in 17.5 μL of nuclease-free water, and 16.75 μL is transferred to a new tube for PCR2.

#### Step 6. PCR2

A second PCR is performed to add unique dual indexes. Representative sequences for P5 and P7 primers are shown in Supplementary Table 1. 16.75 μL of purified PCR1 product is mixed with with dNTPs (0.5 μL, 10 mM stock), P5 and P7 primers (1.25 μL each, 10 μM stock), Phusion High Fidelity proofreading DNA polymerase (0.25 μL Thermofisher), and Phusion HF buffer (5 μL, 5 × stock). For PCR2, initial denaturation was at 98°C for 3 minutes, followed by PCR cycles (98°C 15 seconds, 63°C 30 seconds and 72°C 40 seconds), and final elongation at 72°C for 5 minutes. For high quality RNA input, 17 cycles of PCR are used, and for low quality RNA input, 20 cycles are used.

#### Step 7. Cleanup of PCR2

A final purification step using CleanNGS beads at 0.7x (17.5 μL) is performed before library quantification and sequencing. The final purified product is diluted in 25 μL of nuclease-free water.

#### Step 8. Sequencing

Libraries are quantified using the Qubit fluorometer (Thermo Fisher), fragment size is assessed using the Agilent Tapestation 4200. Successful libraries yield a single broad peak with fragment size between 350-400bp. Libraries are pooled and sequenced on an Illumina MiniSeq or NextSeq (150bp paired end reads) with a 15% PhiX spike-in.

### 5’-RACE library preparation and sequencing

5’-RACE TCRseq libraries were prepared and sequenced as previously described^5^.

### Immunoverse TCRseq library preparation and sequencing

We used the Immunoverse-HS TCR beta/gamma Kit, for Illumina as per manufacturer’s instructions. Libraries were sequenced as described for FUME-TCRseq, with the same target depth.

### ImmunoSEQ library preparation and sequencing

DNA-based sequencing of the TCRβ chain was done using the ImmunoSEQ kit (Adaptive Biotechnologies) according to the manufacturer’s recommendations. The first round of PCR was carried out using the ImmunoSEQ proprietary PCR primer mix (32 μL per sample containing 25 uL of QIAGEN 2× Multiplex PCR Master Mix, 5μL of QIAGEN 5x Q-solution and 2 μL of primer mix). A positive control reaction, provided in the kit, and a negative control reaction were included with each sample batch. PCR cycling parameters were: heated lid (105 °C), 95 °C 5 min denaturation step, followed by 21 cycles of 94 °C, 30 s denaturation, 65 °C, 75 s annealing, and 72 °C, 40 s extension; followed by 72 °C, 10 min final extension, then hold at 4 °C. Amplified libraries were diluted using the DNA suspension buffer (30 μL) provided. A second round of PCR was performed to generate uniquely barcoded sequencing libraries using the barcode primer plate included in the kit (17μL of working mix which include 12.5μL QIAGEN 2x Multiplex PCR master mix, 2.5μL QIAGEN 5x Q-solution and 2μL of QIAGEN RNase-free water; 4 μL of primers from the provided barcode plates and 4 μL of the first PCR product). Second PCR cycling parameters were: heated lid (105 °C), 95 °C

15 min denaturation step, followed by 21 cycles of 94 °C, 30 s denaturation, 68 °C, 40 s annealing, and 72 °C, 60 s extension; followed by 72 °C, 10 min final extension, then hold at 12 °C. The quality of the libraries was assessed using Agilent 4200 TapeStation High Sensitivity D1000 ScreenTape. Samples were pooled volumetrically and purified using MAGBIO HighPrep PCR beads (1×). The final pool was quantitated using a Kapa Library Quantification kit for Illumina. Libraries were sequenced on the Illumina MiSeq System following the manufacturer’s instructions and using MiSeq Reagent Kit v3 (150-cycle) Single-Read. A total of 168 sequencing cycles were performed (Read 1 156 cycles, Read 2 12 cycles), as recommended in the protocol, adding 5% PhiX.

### TCR analysis

For FUME-TCRseq, forward and reverse reads were merged using Vsearch^11^. Processing of FUME-TCRseq and ligation-based TCR sequencing used a previously described suite of Python scripts^5,12^ available at https://github.com/innate2adaptive/Decombinator.

The ImmunoSEQ platform was used for TCR identification and CDR3 extraction of DNA samples sequenced using the ImmunoSEQ kit (Adaptive Biotechnologies). Further analysis of the TCR repertoires was performed in R using the Immunarch package^13^.

For libraries sequenced with the Immunoverse-HS TCR beta/gamma Kit, the Archer Analysis platform was used for TCR identification and CDR3 extraction.

### BaseScope point mutation detection

The detection of KRAS G12C and PIK3CA mutations on FFPE sections was performed using the BaseScope assay (Advanced Cell Diagnostics) as previously described^7^.

## Acknowledgments

This work was funded by Cancer Research UK (A19771, A-MB and TAG) and the Rosetrees Trust and the NIHR UCLH BRC (BC). PN, KH, SB, AM acknowledge funding from Cancer Research UK ICR/RM RadNet Centre of Excellence (C7224/A28724) and the RM/ICR NIHR Biomedical Research Centre. PN acknowledges funding from CRIS Cancer Foundation.

The authors thank Barts Cancer Institute Histopathology Core Facility and The Institute of Cancer Research Histopathology Core Facility for expert processing and sectioning of FFPE blocks.

## Competing interests

Authors A-MB, BC and TAG are listed as co-inventors on patent application GB2305655.9 which relates to the methodological development of FUME-TCRseq. All other authors have no competing interests.

---

**Supplementary Table 1.**
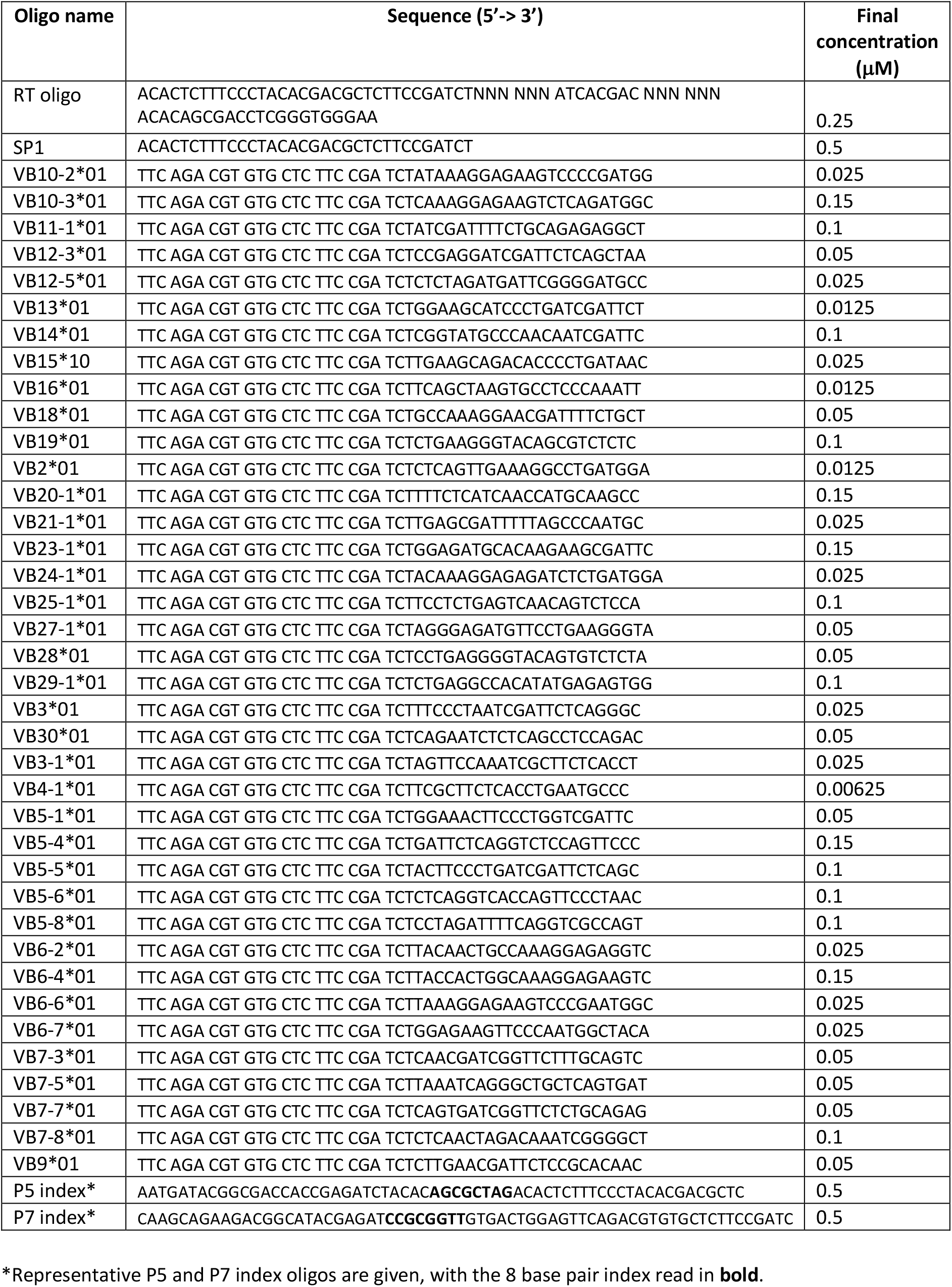
Oligonucleotide details.

**Supplementary Table 2.** Summary of samples and TCR sequencing data.

|  | Type | Figure | RIN | Method | Number of reads input | Number of reads decombined | Number of reads decombined (subsampling) | Total clonotypes | Productive clonotypes | Non productive clonotypes |
| --- | --- | --- | --- | --- | --- | --- | --- | --- | --- | --- |
| Blood | FF | 2 | 7.9 | RACE | 877926 | 538942 | N/A | 18359 | 15512 | 2847 |
| Blood | FF | 2 | 7.9 | FUME-TCRseq | 1353138 | 1325926 | N/A | 143425 | 137432 | 5993 |
| Blood_ subsample | FF | 2 | 7.9 | FUME-TCRseq | 1353138 | 1325926 | 538942 | 118869 | 113981 | 4888 |
| IBD polyp | FF | 2 | 7.8 | FUME-TCRseq | 932201 | 893031 | N/A | 2273 | 2100 | 173 |
| IBD polyp | FF | 2 | 7.8 | FUME-TCRseq | 938189 | 904316 | N/A | 1936 | 1787 | 149 |
| C537 WT | FFPE | 3+5 | 1 | FUME-TCRseq | 922619 | 594987 | N/A | 961 | 727 | 234 |
| C537 MUT | FFPE | 3+5 | 1 | FUME-TCRseq | 1101308 | 796627 | N/A | 1014 | 810 | 204 |
| C539 WT | FFPE | 3 | 2.3 | FUME-TCRseq | 37234 | 32010 | N/A | 129 | 110 | 19 |
| C539 WT_ subsample | FFPE | 3 | 2.3 | FUME-TCRseq | 37234 | 32010 | 18482 | 125 | 108 | 17 |
| C539 MUT | FFPE | 3 | 2.3 | FUME-TCRseq | 26053 | 18482 | N/A | 58 | 48 | 10 |
| Adenoma | FFPE | 4 | 2.3 | FUME-TCRseq | 32224 | 23039 | N/A | 233 | 184 | 49 |
| Carcinoma | FFPE | 4 | 2.3 | FUME-TCRseq | 36389 | 22886 | N/A | 235 | 192 | 43 |
| C537 WT | FFPE | 5 | 1 | Immunoverse | 845042 | N/A | N/A | 478 | 356 | 122 |
| C537 MUT | FFPE | 5 | 1 | Immunoverse | 873394 | N/A | N/A | 317 | 232 | 85 |
| HN1 | FFPE | 6 | 2.6 | FUME-TCRseq | 117174 | 93208 | N/A | 12632 | 9166 | 2466 |
| HN2 | FFPE | 6 | 2.4 | FUME-TCRseq | 113437 | 95977 | N/A | 5805 | 4530 | 1275 |
| HN3 | FFPE | 6 | 2.6 | FUME-TCRseq | 142133 | 111457 | N/A | 18348 | 14588 | 3760 |
| HN1 | FFPE | 6 | 2.6 | ImmunoSEQ | unknown | 65760 | N/A | 3991 | 3251 | 740 |
| HN2 | FFPE | 6 | 2.4 | ImmunoSEQ | unknown | 6874 | N/A | 26862 | 20630 | 6232 |
| HN3 | FFPE | 6 | 2.6 | ImmunoSEQ | unknown | 154701 | N/A | 90070 | 74297 | 15773 |

## Notes

### Summary of Updates

Addition of Supplementary Table 2. In FFPE samples, the filtering of variants represented by only one UMI was removed. Title and text changes to clarify method novelty and utility.

